# Drug Repurposing using consilience of Knowledge Graph Completion methods

**DOI:** 10.1101/2023.05.12.540594

**Authors:** Roger Tu, Meghamala Sinha, Carolina González, Eric Hu, Shehzaad Dhuliawala, Andrew McCallum, Andrew I. Su

## Abstract

While link prediction methods in knowledge graphs have been increasingly utilized to locate potential associations between compounds and diseases, they suffer from lack of sufficient evidence to explain why a drug and a disease may be indicated. This is especially true for knowledge graph embedding (KGE) based methods where a drug-disease indication is linked only by information gleaned from a vector representation. Complementary pathwalking algorithms can increase the confidence of drug repurposing candidates by traversing a knowledge graph. However, these methods heavily weigh the relatedness of drugs, through their targets, pharmacology or shared diseases. Furthermore, these methods can rely on arbitrarily extracted paths as evidence of a compound to disease indication and lack the ability to make predictions on rare diseases. In this paper, we evaluate seven link prediction methods on a vast biomedical knowledge graph for drug repurposing. We follow the principle of consilience, and combine the reasoning paths and predictions provided by path-based reasoning approaches with those of KGE methods to identify putative drug repurposing indications. Finally, we highlight the utility of our approach through a potential repurposing indication.

## Introduction

The estimated cost to bring a drug to market increased from $802 million to $2.7 billion between 2003 and 2013 due to drug attrition, longer development timelines and changing regulatory requirements, adjusting for inflation [1]. Drug repurposing, the process of identifying a new indication for an advanced clinical compound or approved drug, has become increasingly more attractive by leveraging prior work characterizing a drug candidate’s safety and efficacy profile; resulting in a concomitant decrease in time to market, risk of failure, and investment costs [2].

An emerging area in computational drug repurposing exploit features in biomedical knowledge graphs, structured network representations of biomedical facts, to uncover potential links between drugs and diseases [3]. Embedding-based knowledge graph completion techniques learn a latent representation of each entity and relation [4–7].

These methods can infer relationships between drugs and diseases without the limitation of traversing a knowledge graph. However, they lack the ability to provide explanations for their predictions. Path-based knowledge graph completion techniques trace links in the graph and produce a logical sequence to identify and support probable drug repurposing candidates. These methods frequently rely on shared entity targets or entity-entity similarities, which limits their ability to offer convincing and interpretable evidence for a repurposing target [8–10]. This limitation is especially apparent in sparsely connected entities, such as rare diseases.

To enhance predictive modeling for drug repurposing, ensemble approaches can be applied to merge characteristics from various models to create a more robust and accurate model [11]. These approaches have shown promising results in integrating embedding-based models to improve predictive performance. For instance, Bang *et al*. generated embeddings using the word2vec skip-gram algorithm on node sequences extracted from random walks with teleports. An ensemble classifier for drug repurposing utilizing gradient boosting on decision trees was then trained using the embedding features [12]. However, this ensemble approach may not fully recapitulate all facets of drug-disease representations since they train a series of weak models on residual features from a single embedding representation.

Alternatively, ensemble approaches can also integrate different types of embedding-based models to boost predictive performance. Islam *et al*. demonstrated that pre-training embedding representations by concatenating three embedding methods allowed them to train a new link prediction model for predicting drug-disease associations [13]. However, this approach requires a separate path tool to extract logical reasoning chains between a drug repurposing candidate and target disease after deriving associations, rather than integrating a path-based knowledge graph completion algorithm directly with the ensemble methodology.

In this paper, we apply an ensemble voting strategy to drug repurposing. Our methodology is inspired by the principle of consilience, that evidence from independent methods can converge to support a conclusion. This approach extends existing ensemble models by incorporating predictions from both path- and embedding-based approaches, rather than relying solely on pre-trained embeddings that guide a downstream ensemble classifier. Each method, which models distinct representations of the graph, can equally contribute to prioritizing drug repurposing candidates. As a result, this strategy addresses both sparsity and evidence challenges associated with path- and embedding-based approaches, respectively. Here, we train seven path- and embedding-based knowledge graph completion methods independently on a large biomedical knowledge graph using approved drug-disease indications and highlight their drug repurposing ranking performance. Next, we employ an ensemble voting strategy that leverages the complementary strengths of embedding- and path-based methods to prioritize predicted results by its average position. We then contrast the performance of our simplistic ensemble approach against each of its parts. Following, we conduct a thorough analysis on how combinations of disparate knowledge graph completion methods affect ranking performance. Finally, we demonstrate the effectiveness and utility of our ensemble approach to prioritize plausible drug repurposing indications through manual curation.

## Materials and methods

### Notation

A knowledge graph (*G*) can be defined as a collection of edges (*E*) composed by the set of all nodes (*N*) and relations (*R*). Each node can be represented as *n* ∈ *N*, and each relation represented as *r* ∈ *R*, where an edge *e* ∈ *E* ⊆ (*N × R × N*) and can be written as (*n*_1_, *r, n*_2_). An example of an edge is (imatinib, *treats*, chronic myelogenous leukemia [CML]), where “imatinib” and “CML” are nodes, and “*treats*” is a relation. A path is a sequence of interconnected edges and can be written as (*n*_1_, *r*_1_, *n*_2_, …, *r*_*i*_, *n*_*i*_), where *i* represent the number of nodes and relations in the path. An example of a path is (imatinib, *inhibits*, ABL, *associated with*, BCR-ABL, *marker of*, CML). A metapath is a path schema that reflects the node and relation types comprising a path or set of paths. Given the aforementioned path, an example of its metapath is (compound, *inhibits*, gene, *associated with*, gene, *marker of*, disease).

### <u>M</u>echanistic Repurposing Network with <u>Ind</u>ications (MIND)

MIND is a knowledge graph that distinguishes regulatory-approved drug indications from drug-disease relationships from other drug-disease relationships. MIND is rooted in MechRepoNet, a knowledge graph reflecting important drug mechanism relationships identified from a curated dataset of biomedical drug mechanisms [14]. By solely training and evaluating models on *indications* instead of the broader *treats* edge connecting drugs and diseases, MIND more precisely replicates the challenge of predicting approved indications compared to MechRepoNet. The *indications* edges were integrated from DrugCentral, a curated resource with regulatory-approved drug indications [15]. In total, MIND consists of 9,652,116 edges, 249,605 nodes, 9 node types and 22 relations. Supplementary Acknowledgments 1 (upper) highlights total node to node and (lower) node to relation counts in MIND as a whole.

### Applied drug repurposing algorithms

In this paper, we utilized and evaluated a variety of knowledge graph completion algorithms for drug repurposing on the MIND knowledge graph. These algorithms, described in further detail below, fall in two classes: knowledge graph embeddings, and path reasoning methods.

### Knowledge graph embeddings

Knowledge graph embedding (KGE) algorithms leverage a score function, *f* (*n*_2_|*n*_1_, *r*), to learn a vector representation of each node and relation in a latent space. In this function, *n*_2_ represents the predicted answer given a query *n*_1_ and *r*. The primary objective is to rank known *n*_2_ answers higher than unknown *n*_2_ nodes. We trained KGE models on MIND to predict drug repurposing candidates with the following algorithms: TransE [4], DistMult [5], ComplEx [16] and RotatE [7]. Each algorithm’s scoring functions are provided in Supplementary Acknowledgments.

### Path reasoning methods

Path reasoning methods leverage and traverse knowledge graph edges to identify potential drug repurposing candidates. Here, we trained Rephetio [14], Case based reasoning (CBR) [17], and probabilistic CBR [18] models on MIND to predict drug repurposing candidates. These methods can be classified into two classes of path-based reasoning methods: Degree Weighted Path Count (Rephetio) and Case-based reasoning. Degree Weighted Path Count is an algorithm from the Rephetio project that penalizes paths traveling through high-degree nodes when calculating metapath (path based on node type) prevalence; metapaths are incorporated into a logistic regression to calculate the expected probability a compound treats a disease [19]. Mayers *et al*. expanded on this approach and incorporated rules based path exclusions and hyperparameter optimization schemes to streamline operation and improve path interpretability for drug repurposing [14].

CBR is an established technique from artificial intelligence modeled after human ability to retrieve and apply prior experience to tackling a new but similar challenge [20–24]. Briefly, CBR models first retrieve similar entities to the query entity, *n*_1_, that have the specified query relation, *r* given a query (*n*_1_, *r*). Next, the set of relations connecting the similar entities to their answer via r are collected and the paths obtained are applied to the query entity *n*_1_ to identify the answer [17]. Probabilistic CBR extends the original CBR method by utilizing probabilistic models to estimate the likelihood a retrieved path is correct given the query relation [18]. CBR approaches represent a promising avenue for drug repurposing, as it leverages solutions (paths) gathered between existing similar drugs and their diseases they treat to identify potential therapies.

### Drug repurposing with consilience

To prioritize drug repurposing candidates, we implemented a basic ensemble voting approach inspired by the consilience principle, which proposes that evidence from unrelated sources converge on solid conclusions. Firstly, we trained seven knowledge graph completion methods independently and measured their ability to correctly rank approved indications over unknown drug-disease links. Next, we utilized and evaluated two simple ensemble strategies to prioritize drug repurposing candidates: intersection and union ensemble (Fig 1). Finally, we validated the plausibility of top predictions following our ensemble strategies by conducting a literature review.

**Fig 1.**
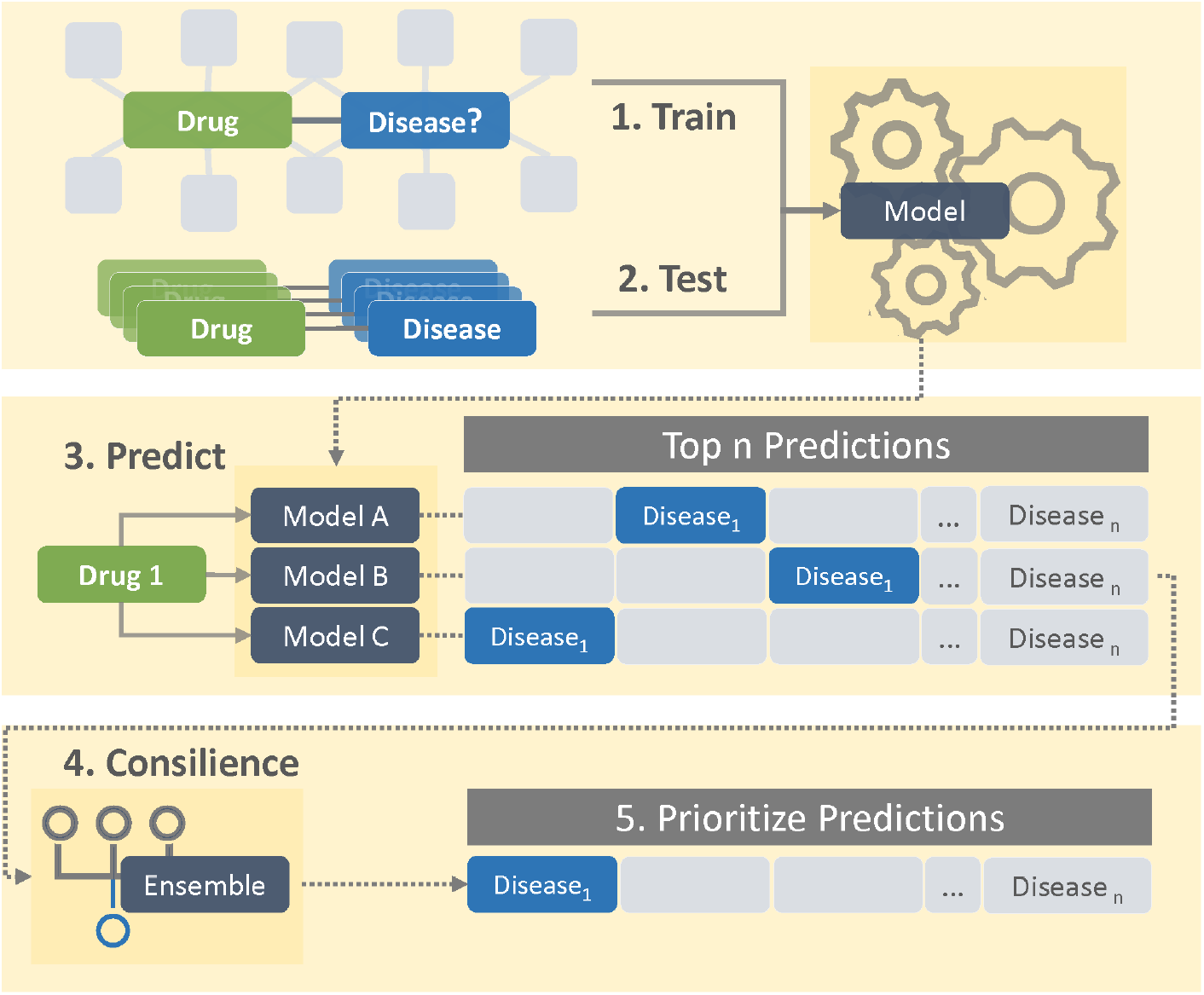
Consilience inspired drug repurposing schematic. 1: Each knowledge graph completion model is trained on the same biomedical knowledge graph and 2: evaluated on approved drug indications (test set). 3: Given a compound, each model predicts diseases the compound may treat. 4: The common predicted disease ranks are collected and scored. 5: Diseases are prioritized to identify drug repurposing candidates.

### Intersection and Union ensemble strategies

For each drug-disease indication in the test set, predictions were made for each knowledge graph completion approach. These predictions can be represented by *P*, where 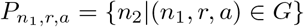 represents the set of predictions for each algorithm *a*, and query (*n*_1_, *r*); *P* can be bounded to keep the first *k* predictions. The predictions were then aggregated into algorithm combinations ranging from two to seven following an intersection or union strategy. An intersection policy is used to filter predictions, ensuring that only those reported by all algorithms in the set of algorithms are retained; the intersection operation can be generalized as:

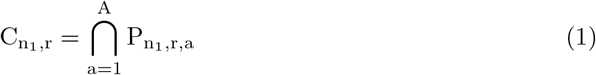

where *A* is the set of algorithms ensembled, *C* is the set of consensus predictions between all models for a given query, *c* is a concept in *C*, and *C* ⊆ *P*. To maintain consistency among algorithms, a union policy stores all predictions made by any algorithm and prioritizes predictions that are unanimously reported by all algorithms in the set. This prioritization is achieved by imposing a penalty on predictions that lack consensus; the union operation can be generalized as:

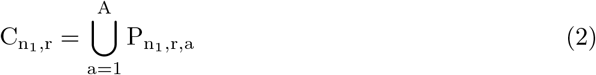

Aggregated predictions were prioritized based on the calculated score. Essentially, the priority score of a prediction is an average value which is penalized with a user defined parameter when the prediction is not forecasted by all the methods in the set. The score function can be represented as:

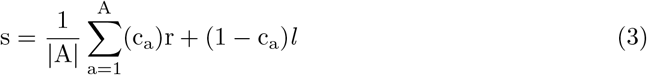

where *c*_*a*_ is a binary value denoting the presence of prediction by an algorithm (1 if present, 0 otherwise), *x* the rank of the prediction, *l* the penalty applied, and |*A*| the number of algorithms combined. Finally, the score function is applied to all predictions for 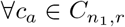 and *C* is indexed in a monotonically increasing order by the score to prioritize consensus predictions. We chose a prediction cutoff *k* = 100, and assigned a penalty *l* as *k* + 1.

Statistical testing was conducted on the rank distributions of approved indications vs not approved predictions grouped by algorithm combination lengths using the Kruskal-Wallace H-test; a non-parametric method of comparing medians of groups. The Mann-Whitney U-test was applied to observe differences between the distributions of approved and unknown drug to disease indications.

### Evaluating plausibility of predicted indications

To evaluate the plausibility of drug repurposing candidates, a literature review was conducted for each drug and its primary disease candidate (first prioritized prediction) from the test set. Every potential indication was classified into three categories: positive, negative, or neutral effect. An indication prediction was labeled “positive” if the literature review suggested that a drug treated or improved the disease condition; these predicted edges have the potential to become drug indications, warrant further study, or are already used clinically off-label. “Negative” labels represent predicted indications that may induce, negatively impact, or worsen the drug’s effect on the disease. Lastly, “neutral” labeled edges have no literature support for the given drug and disease prediction or have been found to neither treat nor modify disease outcomes. The curation results are available in the Supplementary Acknowledgments.

### Model hyperparameter optimization, training, and evaluation

Each embedding- and path-based knowledge graph completion method hyperparameters were tuned over MIND train and validation splits and evaluated on the test split utilizing a compound-to-disease indication train/test/validation split of 80/10/10%, respectively. Algorithms were evaluated using hits at *k*, the success rate of identifying the correct answer among the top *k* predictions, and mean reciprocal rank (MRR), the average of the reciprocal of the positional rank for all correct answers. KGE and CBR hyperparameters were optimized using *Optuna* software [25]. Rephetio was optimized independently through its own hyperparameter optimization pipeline. Hyperparameter optimization selected parameter values can be found at Supplementary Acknowledgments and Acknowledgments, respectively.

## Results

In this section, we applied several knowledge graph completion algorithms on the MIND dataset to identify drug repurposing candidates using our consilience-inspired ensemble method. First, we trained and evaluated the performance of each knowledge graph completion algorithm independently on DrugCentral indications in MIND and compared the parts against our ensemble approach. Next, we studied the ability of two ensemble policies, union and intersection, to rank true indications and unknown edge predictions. Following, we investigated the plausibility of our putative top drug repurposing candidates indications by gathering supporting evidence through manual review of the literature. Finally, we explored the mechanism of action of a potential drug repurposing indication.

### Knowledge graph indication prediction performance comparison

In this study, we assessed the effectiveness of individual and ensemble methods in predicting approved drug-disease indications using the MIND dataset. We conducted training and testing to evaluate the performance of each approach. Among the path traversal methods, probCBR exhibited the best performance, achieving a mean reciprocal rank (MRR) of 0.2557. TransE emerged as the highest-performing embedding-based approach, with an MRR of 0.1601. Turning to ensemble methods, the application of the intersection policy across all seven knowledge graph completion methods outperformed each individual algorithm and yielded the highest overall MRR of 0.9792. Conversely, the union policy resulted in a moderate MRR of 0.1613, on par with TransE and weaker performing than probCBR and the intersection approach. Table 1 presents a summary of the prediction performance for each algorithm individually and when combined in ensembles.

**Table 1.**
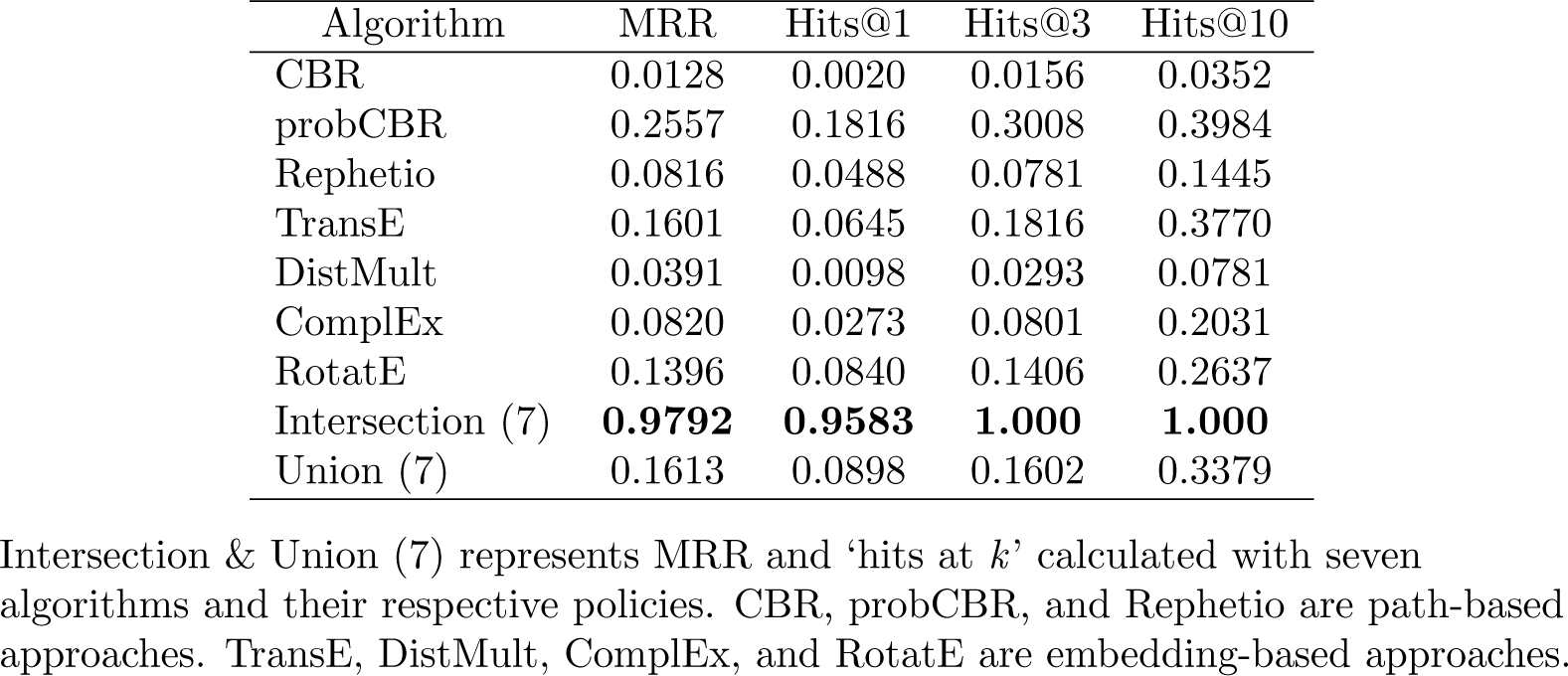
Individual and ensemble methods prediction performances on MIND test set.

| Algorithm | MRR | Hits@1 | Hits@3 | Hits@10 |
| --- | --- | --- | --- | --- |
| CBR | 0.0128 | 0.0020 | 0.0156 | 0.0352 |
| probCBR | 0.2557 | 0.1816 | 0.3008 | 0.3984 |
| Rephetio | 0.0816 | 0.0488 | 0.0781 | 0.1445 |
| TransE | 0.1601 | 0.0645 | 0.1816 | 0.3770 |
| DistMult | 0.0391 | 0.0098 | 0.0293 | 0.0781 |
| ComplEx | 0.0820 | 0.0273 | 0.0801 | 0.2031 |
| RotatE | 0.1396 | 0.0840 | 0.1406 | 0.2637 |
| Intersection (7) | **0.9792** | **0.9583** | **1.000** | **1.000** |
| Union (7) | 0.1613 | 0.0898 | 0.1602 | 0.3379 |

Intersection & Union (7) represents MRR and ‘hits at $k$ ’ calculated with seven algorithms and their respective policies. CBR, probCBR, and Rephetio are path-based approaches. TransE, DistMult, ComplEx, and RotatE are embedding-based approaches.

### Ensemble policy effect on predicted indications

#### Ensemble mean reciprocal rank performance and counts

Next, we investigated ensemble policy effects on ranking approved drug-disease indications and drug repurposing candidate counts. Fig 2a shows the changes in MRR prediction performance and Fig 2b highlights the candidate prediction counts as the number of algorithms included in the ensemble predictor increases. For a given ensemble predictor size in Fig 2, all possible combinations of algorithms are averaged.

**Fig 2.**
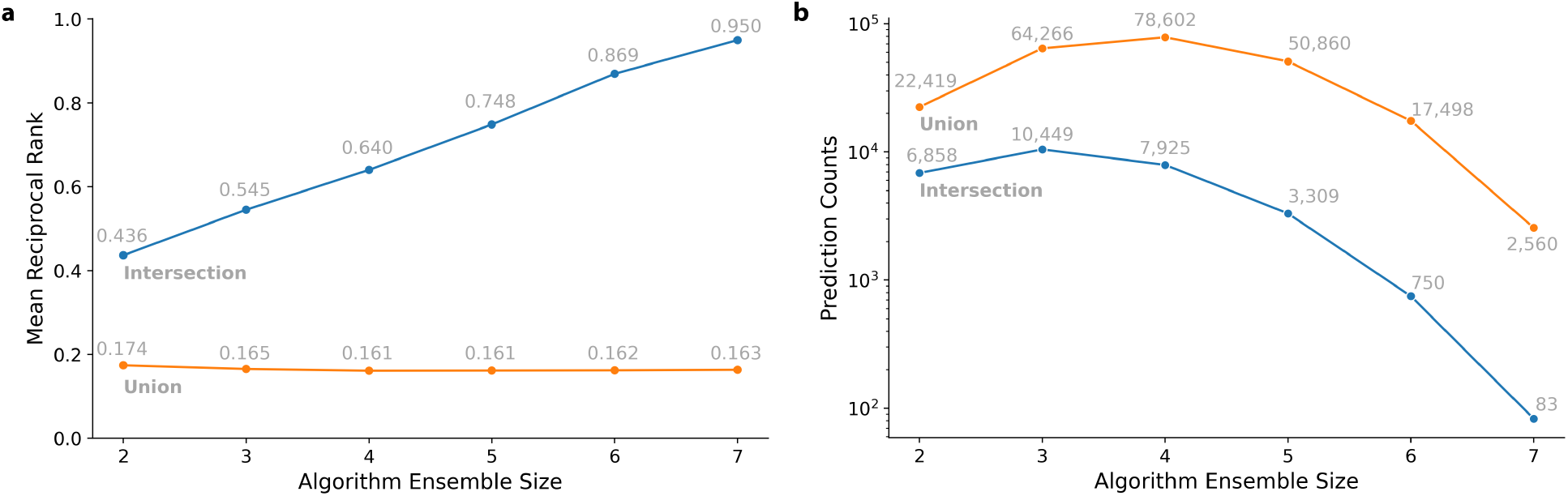
Ensemble ranking performance & count statistics for intersection & union policies. **a:** Approved indication prediction MRR with respect to algorithm ensemble size. **b:** Indication counts with respect to algorithm ensemble size; the y-axis is log scaled. Blue and orange lines denote intersection and union strategy, respectively.

Following an intersection policy (Fig 2a, blue line), we observed a steady increase in MRR with increasing algorithm ensemble size. This approach improved MRR performance from 0.436 to 0.950 when algorithm ensemble count increased from two to seven, respectively. However, the intersection policy’s prospective indications dropped sharply as the algorithm ensemble size increased (Fig 2b, blue line); only 83 candidate indications remained when algorithm combination length was seven.

Based on a union strategy (Fig 2a, orange line), the MRR value slightly decreased with increasing algorithm ensemble size. Instead, the MRR performance experienced a slight decrease from 0.174 to 0.163 as the ensemble count increased from two to seven, respectively. In contrast to the intersection policy, the candidate indication counts (Fig 2b, orange line) exhibited a concave pattern as more algorithms were ensembled. Consequently, the union policy candidate indication count started at 22,419, reached a peak of 78,602, and concluded at 2,560. This is likely not only because there are more diverse predictions with a union approach, but also because there are many different combinations that can be included in an ensemble.

#### Ensemble ranking performance of approved indications

We compare the ranking performance of our consilience-inspired approaches using known drug-disease indications. Fig 3 demonstrates the rank of approved indications (positive) with respect to non-indications (negative) rank distributions for the intersection and the union policy across varying algorithm ensemble sizes.

**Fig 3.**
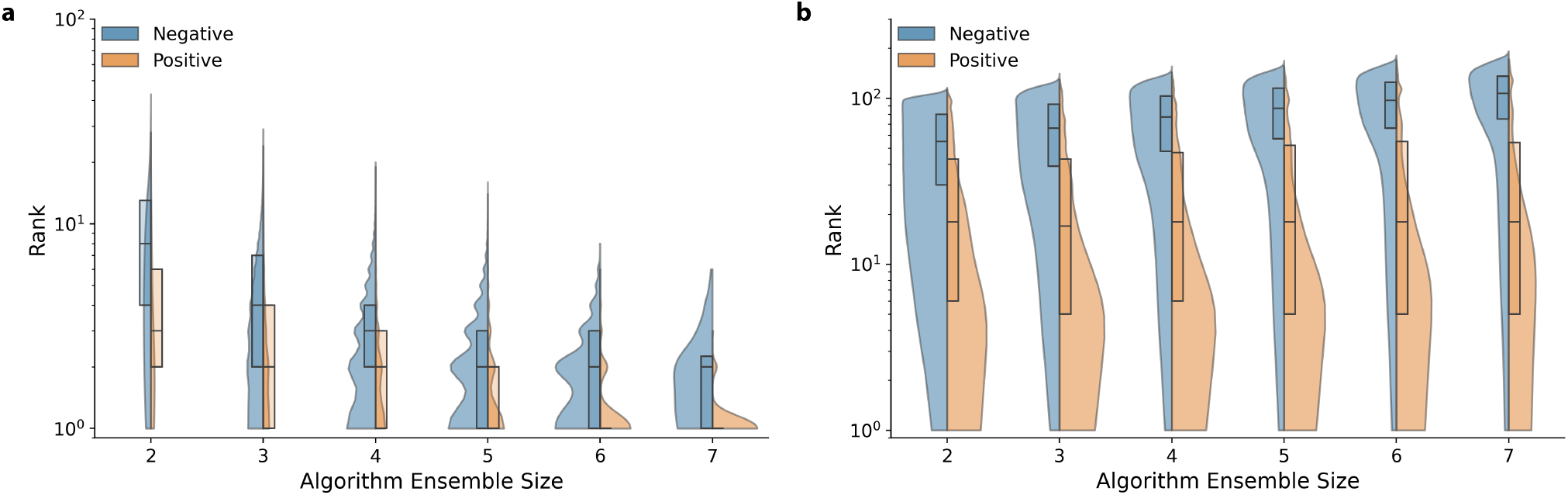
Intersection & union policy effect on indication ranking performance. Positive (approved drug-disease indications) and negative (non-indications) ensemble prediction distributions were visualized. **a:** Intersection policy effect on ensemble ranking distribution of positive and negative sets. **b:** Union policy effect on ensemble ranking distribution of positive and negative sets. X-axis represents ensemble combinations between CBR, probCBR, Rephetio, TransE, DistMult, ComplEx and RotatE of sizes two through seven. Boxplot distributions highlight the first, second and third quartiles for Positive and Negative prediction ranks. The distribution between Positive and Negative sets are statistically significantly different regardless of algorithm combination length and ensemble policy. Violin plots highlight the kernel density estimate for each policy’s Positive and Negative rank distribution; the y-axis is log scaled.

Subject to an intersection policy, Fig 3a highlights strong ranking performance among both “positive” and “negative” groups, with approved indications ranking better than non-indication predictions. The median indication rank was 1 with 83 counts, whereas the median “negative” group rank was 2 with 68 counts at an ensemble size of seven. Applying the Kruskal-Wallis H test showed at least one median was statistically different from the others; subsequent statistical testing using the Mann-Whitney U test demonstrated the positive and negative distributions for each ensemble size were statistically significant (*p <* 0.001).

Following a union policy, the median rank of the “positive” and “negative” groups was 18 with 2,560 counts, and 107 with 180,231 counts with an ensemble size of seven, respectively (Fig 3b). The Kruskal-Wallis H-test indicated at least one distribution median was statistically different from the others. Similar to the intersection policy, subsequent statistical testing using the Mann-Whitney U-test showed that each pair of positive and negative distributions for each ensemble size was statistically significant (*p <* 0.001). Supplementary Acknowledgments and Acknowledgments show the Kruskal-Wallis H-test and Mann-Whitney U-test statistics and p-value, respectively.

Overall, the findings indicate that both the intersection and union policies significantly enhance the ranking performance for approved indications. However, the intersection policy results in fewer potential drug repurposing candidates as it relies on collaborative agreement between all algorithms. It is noteworthy that while the ranking performance of known indications is inferior to that of the intersection policy, the union policy effectively prioritizes known indications over non-indications.

#### Inference evaluation through literature curation

To evaluate the inference performance of our ensemble approach compared to its parts, we conducted a literature curation exercise on 25 randomly selected drugs and their predicted diseases. We chose to curate predictions from the union over the intersection policy as it represents a more realistic prediction set for estimating ensemble predictive ability. Each sampled drug was categorized as having a positive, negative, or neutral impact on its predicted disease. Based on our assessment, the union ensemble policy demonstrated superior performance compared to each algorithm individually with 80%, 4% and 16% of the sampled drugs classified as positively, negatively, and not affecting the predicted diseases, respectively. Among path- and embedding-based approaches, Rephetio and TransE had the most negatively and positively classified predictions, respectively. Despite outperforming other methods on the test set, probCBR’s performance in our curation exercise was unremarkable. The complete curation results are provided in the Supplementary Acknowledgments. Table 2 summarizes the curation.

**Table 2.** Ensemble inference performance results comparison.

|  | Union (7) | CBR | probCBR | Rephetio | TransE | DistMult | ComplEx | RotatE |
| --- | --- | --- | --- | --- | --- | --- | --- | --- |
| Positive | 80% (20) | 28% (7) | 48% (12) | 48% (12) | 72% (18) | 64% (16) | 68% (17) | 48% (12) |
| Neutral | 16% (4) | 60% (15) | 48% (12) | 32% (8) | 24% (6) | 36% (9) | 28% (7) | 48% (12) |
| Negative | 4%(1) | 12% (3) | 4% (1) | 20% (5) | 4%(1) | 0%(0) | 4% (1) | 4% (1) |

Through literature curation, the top predicted disease for each drug for each given algorithm was categorized into three groups by the effect the drug has on its predicted indication; the groups are positive, neutral, and negative effects. An indication was categorized as positive if a drug improved the predicted disease outcome, negative if a drug exacerbated the predicted disease, or neutral if a predicted disease was neither affected nor associated with a drug.

#### Case study: Sotalol hydrochloride as a potential treatment for hypertension

Among the most confident predictions made by our consilience inspired approach was the use of sotalol hydrochloride (sotalol) to treat hypertension. This predicted indication was made by all seven algorithms at ranks ranging from 1 to 86 as seen in Supplementary Acknowledgments. Sotalol, an atypical beta blocker, is approved to treat arrhythmias like atrial fibrillation and ventricular tachycardia. While sotalol has not been approved for hypertension, various clinical studies have demonstrated its efficacy in controlling hypertension independently and as an adjunctive therapy with thiazides [26–28]. Utilizing our computational repurposing approach, we propose three mechanisms by which sotalol may moderate hypertension (a condition characterized by increased arterial blood pressure) through potassium channel activity, ADRB1 and FNDC4. Each prospective mechanism retrieved were the top ranked paths by each path reasoning method and is illustrated in Fig 4.

**Fig 4.**
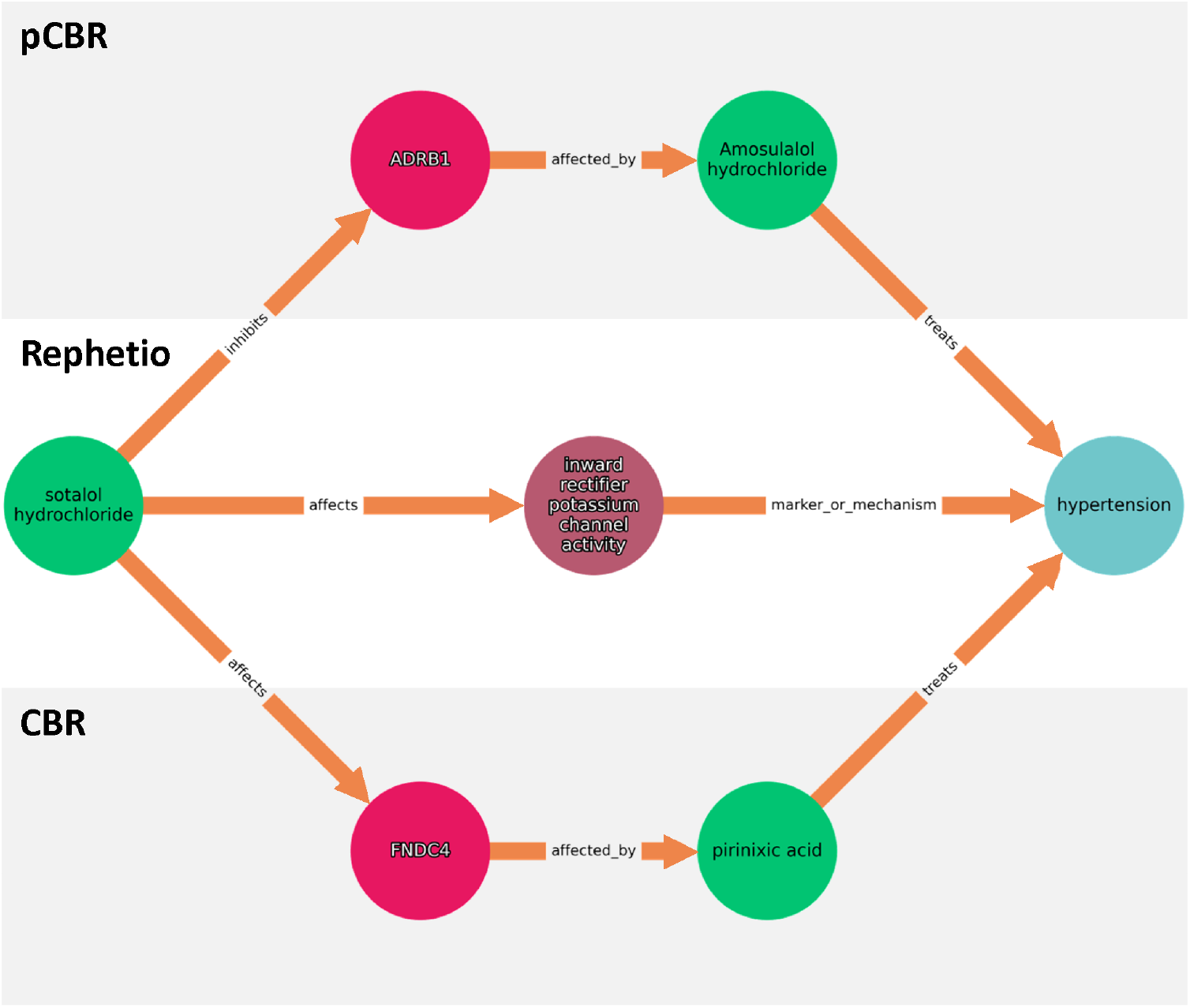
Three prospective mechanisms that sotalol hydrochloride may treat hypertension. Each path shown was the first extracted path suggested by probCBR, Rephetio and CBR for the putative indication. Green, purple, pink and blue circles represent compounds, biological processes, genes & proteins, and diseases, respectively. Orange arrows in between nodes describe node relationships.

A canonical avenue sotalol may manage hypertension is through inhibition of ADRB1, a *β*-adrenergic receptors. ADRB1 inhibition mediates adrenaline’s effect on the heart, resulting in concomitant decreased blood pressure and cardiac output [29]. As sotalol and amosulalol both directly block ADRB1, a protein and gene associated with hypertension, and amosulalol has anti-hypertensive effects, it is reasonable that sotalol also has anti-hypertensive effects [30, 31].

Another approach sotalol may modulate hypertension is through potassium channel activity [32]. While potassium channel blockers are associated with exacerbating blood pressure as it prevents the outflow of potassium ions, it is plausible that sotalol’s inhibition of inward-rectifier potassium channels (repolarization) induces vasodilation and as a result, decreases blood pressure [33–35].

Finally, sotalol may manage hypertension through upregulating the expression of FNDC4, an anti-inflammatory factor [36]. FNDC4 belongs to the fibronectin type III domain-containing protein family, and is highly homologous to irisin, a myokine derived from the proteolytic cleavage of FNDC5 [37, 38]. Deletion and overexpression of irisin has been shown to exacerbate and ameliorate cardiac hypertrophy in rats with hypertension, respectively [39]. As sotalol and pirinixic acid treatment both upregulate irisin expression, and pirinixic acid has hypertension mediating effects, it is probable that sotalol also exhibits anti-hypertensive properties [40, 41].

## Discussion

In this work, we apply a consilience-based, ensemble approach on seven algorithms and demonstrate an enhanced ability to predict approved and putative indications. By combining path-based and KGE algorithms, our method synergistically bolsters path-based and KGE inspired drug repurposing performance through increased resilience to missing edges and reinforced prospective drug repurposing indications with reasoning chains. Our strategy not only validates KGE derived predictions, but it also allows path-based predictions to prioritize diseases with minimal similar cases in the graph.

Our selection of seven knowledge graph completion approaches was driven by several considerations. For KGE approaches, we began with TransE due to its conceptual simplicity and widespread use. Recognizing that relying on a single embedding-based method would not provide a comprehensive representation of class performance, we explored other seminal works that utilized embeddings for knowledge graph completion. Conveniently, Sun et al had implemented TransE along with DistMult, ComplEx, and RotatE, which we applied for comparative purposes. In terms of path-based approaches, which provide reasoning chains to support drug repurposing candidates, we drew upon our prior work involving Rephetio. To diversify our methodologies, we investigated the application of CBR given its intuitive appeal. Regardless of the specific underlying algorithms, we believe that our ensemble approach has the potential to prioritize drug repurposing candidates effectively.

Although our method viably identifies potential repurposing candidates, one challenge in our implementation is the restrictive nature of the intersection policy. As the number of algorithms applied for consilience increases, the fewer putative indications remain (as illustrated in Figure 2). This issue can be ostensibly mediated by either increasing the number of total predictions made per indication and/or increasing the number of compounds used for inference. Expanding our study limits for each algorithm’s predictions (from 100 to the top 1000 ranked diseases and/or inference compounds from 500 to 1000), barring computational limits, could be considered. An additional avenue to address the intersection policy’s restrictive properties is through the union policy; instead of eliminating popular but nonunanimous predictions, the rank is padded with a tunable user specified rank. This approach preserves intersecting prospective indications and penalizes partially-intersected candidates during the re-ranking step. Notably our study deploying a union ensemble strategy demonstrated a decrease in approved indication performance. This is because the union policy trades precision for breadth of results returned, and can be more useful when combining more methods together in an ensemble.

Another challenge to our approach is the equal weighting of each algorithm’s predictions in its irrespective of prior observed performance on a dataset. Weighting each algorithm’s contributions by the predicted indication rank or by incorporating an algorithm’s cross validation performance into a logistic regression would potentially improve the prediction efficacy of the algorithm. These modifications would further fuel the addition of diverse algorithms into our implementation.

Finally, our utilization of path-based methods does not mitigate similarity based drug-disease associations, even when the results are filtered by those that also occur in KGE approaches. For example, the pathway identified by our approach in the between sotalol hydrochloride and hypertension, traverses through ADRB1, a commonly shared target by amosulalol and sotalol, in order to treat hypertension. While similarity based paths correctly identify analogous family compounds (sotalol and amosulalol are both beta blockers), adding simple path similarity filters or penalties to the path retrieval mechanism would likely improve both the path retrieval algorithm and our own approach.

Regardless of variation of the consilience strategy applied, our results demonstrate that even with a naive approach, we were able to identify putative drug repurposing candidates that are supported by existing literature. Our strategy synergizes the advantages of both KGE and path-based algorithms by blending their combined predictive power and strengths. Through manual literature curation, we demonstrate that our method is more likely to identify a drug repurposing indication than when utilizing each algorithm alone. Moreover, our method enables human interpretable reasoning chains derived from path-based approaches to support a putative compound to disease indication that would otherwise not be present with KGEs alone.

## Acknowledgments

The authors would like to thank Dr. Laura Hughes for visualization feedback, and Dr. Chunlei Wu, Dr. Jean-Christophe Ducom and Dr. Rajarshi Das for computational support. This work was supported by the National Institute of Health (R01 AG066750).

## References

1. DiMasi JA, Grabowski HG, Hansen RW. Innovation in the pharmaceutical industry: New estimates of R&D costs. Journal of Health Economics. 2016 May;47:20–33. Available from: https://linkinghub.elsevier.com/retrieve/pii/S0167629616000291.

2. Ashburn TT, Thor KB. Drug repositioning: identifying and developing new uses for existing drugs. Nature Reviews Drug Discovery. 2004 Aug;3(8):673–83. Available from: http://www.nature.com/articles/nrd1468.

3. Pushpakom S, Iorio F, Eyers PA, Escott KJ, Hopper S, Wells A, et al. Drug repurposing: progress, challenges and recommendations. Nature Reviews Drug Discovery. 2019 Jan;18(1):41–58. Available from: http://www.nature.com/articles/nrd.2018.168.

4. Bordes A, Usunier N, Garcia-Durán A, Weston J, Yakhnenko O. Translating embeddings for modeling multi-relational data. In: Proceedings of the 26th International Conference on Neural Information Processing Systems - Volume 2. NIPS’13. Red Hook, NY, USA: pnCurran Associates Inc.; 2013. p. 2787–95.

5. Yang B, Yih Wt, He X, Gao J, Deng L. Embedding Entities and Relations for Learning and Inference in Knowledge Bases. arXiv; 2015. 1412.6575 [cs]. Available from: http://arxiv.org/abs/1412.6575.

6. Toutanova K, Lin V, Yih Wt, Poon H, Quirk C. Compositional Learning of Embeddings for Relation Paths in Knowledge Base and Text. In: Proceedings of the 54th Annual Meeting of the Association for Computational Linguistics (Volume 1: Long Papers). Berlin, Germany: Association for Computational Linguistics; 2016. p. 1434–44. Available from: http://aclweb.org/anthology/P16-1136.

7. Sun Z, Deng ZH, Nie JY, Tang J. RotatE: Knowledge Graph Embedding by Relational Rotation in Complex Space. arXiv. 2019;abs/1902.10197:18. Available from: http://arxiv.org/abs/1902.10197.

8. Li J, Lu Z. A new method for computational drug repositioning using drug pairwise similarity. In: 2012 IEEE International Conference on Bioinformatics and Biomedicine. Philadelphia, PA, USA: IEEE; 2012. p. 1–4. Available from: http://ieeexplore.ieee.org/document/6392722/.

9. Luo H, Wang J, Li M, Luo J, Peng X, Wu FX, et al. Drug repositioning based on comprehensive similarity measures and Bi-Random walk algorithm. Bioinformatics. 2016 Sep;32(17):2664–71. Available from: https://academic.oup.com/bioinformatics/article-lookup/doi/10.1093/bioinformatics/btw228.

10. Luo H, Li M, Wang S, Liu Q, Li Y, Wang J. Computational drug repositioning using low-rank matrix approximation and randomized algorithms. Bioinformatics. 2018 Jun;34(11):1904–12. Available from: https://academic.oup.com/bioinformatics/article/34/11/1904/4820334.

11. Mahajan P, Uddin S, Hajati F, Moni MA. Ensemble Learning for Disease Prediction: A Review. Healthcare. 2023 Jan;11(12):1808. Number: 12 Publisher: Multidisciplinary Digital Publishing Institute. Available from: https://www.mdpi.com/2227-9032/11/12/1808.

12. Bang D, Lim S, Lee S, Kim S. Biomedical knowledge graph learning for drug repurposing by extending guilt-by-association to multiple layers. Nature Communications. 2023 Jun;14(1):3570. Publisher: Nature Publishing Group. Available from: https://www.nature.com/articles/s41467-023-39301-y.

13. Islam MK, Amaya-Ramirez D, Maigret B, Devignes MD, Aridhi S, Smäil-Tabbone M. Molecular-evaluated and explainable drug repurposing for COVID-19 using ensemble knowledge graph embedding. Scientific Reports. 2023 Mar;13(1):3643. Publisher: Nature Publishing Group. Available from: https://www.nature.com/articles/s41598-023-30095-z.

14. Mayers M, Tu R, Steinecke D, Li TS, Queralt-Rosinach N, Su AI. Design and application of a knowledge network for automatic prioritization of drug mechanisms. Bioinformatics (Oxford, England). 2022 May;38(10):2880–91.

15. Ursu O, Holmes J, Knockel J, Bologa CG, Yang JJ, Mathias SL, et al. DrugCentral: online drug compendium. Nucleic Acids Research. 2017 Jan;45(D1):D932–9. Available from: 10.1093/nar/gkw993.

16. Trouillon T, Welbl J, Riedel S, Gaussier E, Bouchard G. Complex embeddings for simple link prediction. In: Proceedings of the 33rd International Conference on International Conference on Machine Learning - Volume 48. ICML’16. New York, NY, USA: JMLR.org; 2016. p. 2071–80.

17. Das R, Godbole A, Dhuliawala S, Zaheer M, McCallum A. A Simple Approach to Case-Based Reasoning in Knowledge Bases. arXiv. 2020;abs/2006.14198. Available from: https://arxiv.org/abs/2006.14198.

18. Das R, Godbole A, Monath N, Zaheer M, McCallum A. Probabilistic Case-based Reasoning for Open-World Knowledge Graph Completion. arXiv; 2020. 2010.03548 [cs]. Available from: http://arxiv.org/abs/2010.03548.

19. Himmelstein DS, Lizee A, Hessler C, Brueggeman L, Chen SL, Hadley D, et al. Systematic integration of biomedical knowledge prioritizes drugs for repurposing. eLife. 2017 Sep;6:e26726. Publisher: eLife Sciences Publications, Ltd. Available from: 10.7554/eLife.26726.

20. Schank RC. Dynamic memory: A theory of reminding and learning in computers and people. cambridge university press; 1983.

21. Kolodner JL. Maintaining organization in a dynamic long-term memory. Cognitive science. 1983;7(4):243–80. ISBN: 0364-0213 Publisher: Elsevier.

22. Rissland EL. Examples in Legal Reasoning: Legal Hypotheticals. In: IJCAI; 1983. p. 90–3.

23. Aamodt A, Plaza E. Case-based reasoning: Foundational issues, methodological variations, and system approaches. AI communications. 1994;7(1):39–59. ISBN: 0921-7126 Publisher: IOS press.

24. Leake DB. CBR in context: the present and future. Case based reasoning experiences-lessons and future experiences. D. Leake. Cambridge, MIT Press; 1996.

25. Akiba T, Sano S, Yanase T, Ohta T, Koyama M. Optuna: A Next-generation Hyperparameter Optimization Framework. arXiv; 2019. 1907.10902 [cs, stat]. Available from: http://arxiv.org/abs/1907.10902.

26. Fillastre JP, Godin M, Joire JE, Gauthier A. [Treatment of hypertension with Sotalol (author’s transl)]. La Semaine Des Hopitaux: Organe Fonde Par l’Association D’enseignement Medical Des Hopitaux De Paris. 1979 Nov;55(39-40):1825-31.

27. Shaw HL. Once daily sotalol in the treatment of hypertension. The Journal of the Royal College of General Practitioners. 1977 Dec;27(185):742–5. Available from: https://www.ncbi.nlm.nih.gov/pmc/articles/PMC2158659/.

28. Jaattela A. The Combination of Sotalol and Hydrochlorothiazide in the Treatment of Hypertension. The Journal of Clinical Pharmacology. 1979;19(8-9):565–70. eprint: 10.1002/j.1552-4604.1979.tb02523.x. Available from: https://onlinelibrary.wiley.com/doi/abs/10.1002/j.1552-4604.1979.tb02523.x.

29. Betrix L, Timour-Chah Q, Lang J, Lakhal M, Faucon G. Protection against ventricular and atrial fibrillation by sotalol. Cardiovascular Research. 1986 May;20(5):358–63. Available from: 10.1093/cvr/20.5.358.

30. Inoue Y, Yaga K, Nishimura M, Okafuji K, Fujii Y, Nagasaka Y, et al. Antihypertensive and metabolic effects of long-term treatment with amosulalol in non-insulin dependent diabetics. Current Medical Research and Opinion. 1992;12(9):564–71.

31. Huang Y, Liu XL, Wen J, Huang LH, Lu Y, Miao RJ, et al. Downregulation of the beta1 adrenergic receptor in the myocardium results in insensitivity to metoprolol and reduces blood pressure in spontaneously hypertensive rats. Molecular Medicine Reports. 2017 Feb;15(2):703–11. Publisher: Spandidos Publications. Available from: 10.3892/mmr.2016.6038.

32. Edvardsson N, Hirsch I, Emanuelsson H, Pontén J, Olsson SB. Sotalol-induced delayed ventricular repolarization in man*. European Heart Journal. 1980 Oct;1(5):335–43. Available from: 10.1093/eurheartj/1.5.335.

33. Thomas D, Karle CA, Kiehn J. The cardiac hERG/IKr potassium channel as pharmacological target: structure, function, regulation, and clinical applications. Current Pharmaceutical Design. 2006;12(18):2271–83.

34. Li C, Yang Y. Advancements in the study of inward rectifying potassium channels on vascular cells. Channels. 2023 dec;17(1):2237303. Available from: https://www.ncbi.nlm.nih.gov/pmc/articles/PMC10355679/.

35. Chen L, Sampson KJ, Kass RS. Cardiac Delayed Rectifier Potassium Channels in Health and Disease. Cardiac electrophysiology clinics. 2016 Jun;8(2):307–22. Available from: https://www.ncbi.nlm.nih.gov/pmc/articles/PMC4893812/.

36. Atienzar F, Gerets H, Dufrane S, Tilmant K, Cornet M, Dhalluin S, et al. Determination of Phospholipidosis Potential Based on Gene Expression Analysis in HepG2 Cells. Toxicological Sciences. 2007 Mar;96(1):101–14. Available from: 10.1093/toxsci/kfl184.

37. Bosma M, Gerling M, Pasto J, Georgiadi A, Graham E, Shilkova O, et al. FNDC4 acts as an anti-inflammatory factor on macrophages and improves colitis in mice. Nature Communications. 2016 Apr;7(1):11314. Number: 1 Publisher: Nature Publishing Group. Available from: https://www.nature.com/articles/ncomms11314.

38. Frühbeck G, Fernández-Quintana B, Paniagua M, Hernández-Pardos AW, Valentí V, Moncada R, et al. FNDC4, a novel adipokine that reduces lipogenesis and promotes fat browning in human visceral adipocytes. Metabolism. 2020 Jul;108:154261. Available from: https://www.sciencedirect.com/science/article/pii/S0026049520301256.

39. Li RL, Wu SS, Wu Y, Wang XX, Chen HY, Xin Jj, et al. Irisin alleviates pressure overload-induced cardiac hypertrophy by inducing protective autophagy via mTOR-independent activation of the AMPK-ULK1 pathway. Journal of Molecular and Cellular Cardiology. 2018 Aug;121:242–55. Available from: https://www.sciencedirect.com/science/article/pii/S0022282818306965.

40. Leibovitz E, Schiffrin EL. PPAR Activation: A New Target for the Treatment of Hypertension. Journal of Cardiovascular Pharmacology. 2007 Aug;50(2):120. Available from: https://journals.lww.com/cardiovascularpharm/Fulltext/2007/08000/Exercise_Capacity_and_Brain_Natruiretic_Peptide_in.4.aspx.

41. Ma Y, Du Y, Yang J, He Q, Wang H, Lin X. Anti-inflammatory Effect of Irisin on LPS-Stimulated Macrophages Through Inhibition of MAPK Pathway. Physiological Research. 2023 Apr;72(2):235–49. Available from: https://www.ncbi.nlm.nih.gov/pmc/articles/PMC10226406/.

