## supplementary for "Drug Repurposing using consilience of Knowledge Graph Completion methods"

### Supporting Information

**S1 Fig. Describes the composition of nodes and edges in the MIND dataset as a whole.** Upper: Heatmap of node type to node type counts. The y-axis and the x-axis correspond to the head and tail node respectively. Lower: Heatmap of Node type (both head and tail) to relation type counts; Heatmap colors are in exponential scale where darker signifies more and lighter less, respectively. Parentheses highlight the node type and relation type counts.

**S1 Table. Scoring Functions for Knowledge Graph Embedding Algorithms.**

| Algorithm | Score Function |
|---|---|
| TransE | $-\|\|n_1 + r - n_2\|\|$ |
| DistMult | $\langle n_1, r, n_2 \rangle$ |
| ComplEx | $Re\langle n_1, r, n_2 \rangle$ |
| RotatE | $-\|\|n_1 \odot r - n_2\|\|$ |

The angle brackets in DistMult and ComplEx denote a trilinear dot product. denotes the Hadamard product (element wise product) between the n1 and r embeddings. In brief, TransE, short for Translation Embedding, was developed in 2013 by Bordes et al., aimed to model relationships as translations in embedding space; where n2 should be close to the embedding of h when added to a vector, r [**?**]. DistMult, modeled relationships using a bilinear diagonal matrix and trilinear dot product as a scoring function [**?**]. ComplEx modified the DistMult model by embedding additional complex vector components but maintained the trilinear dot product as a scoring function [**?**]. Finally, RotatE modeled each relation as an element-wise rotation in complex vector spaces [**?**].

**S1 File. Curation result of the Union ensemble approach.** 25 randomly selected non-indication predictions were manually curated through a literature search for the union ensemble approach and each of its parts. Each compound was classified as having a "Positive", "Neutral", or "Negative" effect on the predicted disease based on criteria outlined in the methods. In total, 200 predictions were made. The results are summarized in Table **??**.

**S2 Table. KGE approaches Optuna optimization hyperparameter results.**

| Algorithm | Optimized MRR | Batch Size | Hidden Dimension Size | Negative Sample Size | Learning rate |
|---|---|---|---|---|---|
| TransE | 0.054154 | 248 | 225 | 124 | 0.002098 |
| DistMult | 0.021927 | 212 | 300 | 88 | 0.000816 |
| ComplEx | 0.032595 | 252 | 250 | 84 | 0.000292 |
| RotatE | 0.068594 | 240 | 100 | 96 | 0.002329 |

**S3 Table. CBR approaches Optuna optimization hyperparameter results.**

| Algorithm | Optimized MRR | Max Num Programs | Max Path Length | Max Branch | K Adj | Number of Paths Collected | Linkage |
|---|---|---|---|---|---|---|---|
| CBR | 0.09672434757569749 | 25 | 3 | 100 | 10 | 1000 | 0 |
| probCBR | 0.20301439879421163 | 1000 | 3 | 100 | 5 | 1000 | 0 |

**S4 Table. Kruskal-Wallis Test statistic and p-values of various ensemble lengths for each ensemble method.**

| Ensemble group | Indication group | statistic | p-value |
|---|---|---|---|
| Intersection | Positive & Negative | 281752298.0 | 0.0 |
| Intersection | Positive | 134.8568 | 2.2155e-27 |
| Intersection | Negative | 3465.5996 | 0.0 |
| Union | Positive & Negative | 1351570.3371 | 0.0 |
| Union | Positive | 3210.1153 | 0.0 |
| Union | Negative | 1053174.30321 | 0.0 |

We report the Kruskal-Wallis test statistic value as well as the corresponding p-value for three groups: Positive, Negative, and Both. The Positive, Negative, and Both groups conduct the Kruskal-Wallis test on algorithm lengths 2 - 7 and only on approved indications, non-indications, and both approved and non-indications, respectively. Kruskal-Wallis H-Test compares the median of non-parametric distributions to identify if there is a statistically significant difference between the provided distributions. A low p-value suggests that at least one of the distributions is different from the others. .

**S5 Table. Mann-Whitney U-Test statistic and p-values of various ensemble lengths for each ensemble method.**

| | Intersection | | Union | |
|---|---|---|---|---|
| length | statistic | p-value | statistic | p-value |
| 2 | 24670494.5 | 0.0 | 13638459952.0 | 0.0 |
| 3 | 281752298.0 | 0.0 | 145208921890.5 | 0.0 |
| 4 | 102138952.5 | 0.0 | 262928904076.0 | 0.0 |
| 5 | 13011720.0 | 0.0 | 128163422278.0 | 0.0 |
| 6 | 524477.5 | 3.5930e-127 | 17175072890.5 | 0.0 |
| 7 | 5063.0 | 5.2950e-17 | 407357083.0 | 0.0 |

We report the Mann-Whitney U-Test statistic value as well as the corresponding p-value for all six pairs of positive (approved indications) and negative (non-indications) distributions for both the intersection and union ensemble approaches. Mann-Whitney U-Test is a non-parametric method to compare whether two distributions are the same by randomly selecting X and Y from each distribution and measuring if the probability of $X > Y$ is the same as $Y > X$. A low p-value suggests that the probability of $X > Y$ is not the same as the probability of $Y > X$.

**S6 Table. Sotalol hydrochloride's respective predicted rank for each link prediction method.**

| Algorithm | Hypertension Rank |
|---|---|
| CBR | 3 |
| probCBR | 3 |
| Rephetio | 1 |
| TransE | 28 |
| DistMult | 86 |
| ComplEx | 27 |
| RotatE | 1 |
